# Differential thermal sensitivity may explain the temporal distribution of foraging activity among different-sized workers in a polymorphic ant species

**DOI:** 10.64898/2025.12.04.692124

**Authors:** Jessica Josefa Sanches, Maud Combe, Ronald Zanetti, Vincent Fourcassié

## Abstract

One of the most stressful factors for insects is increasing temperature because of the risk of potentially fatal dehydration linked to their small size. We used respirometry to study the effect of both temperature and body mass on water loss and metabolic rate in individual workers of the polymorphic ant species *Messor barbarus.* As expected, we found that large ants exposed to increasing temperatures have a lower rate of water loss than small ants and that their mass-specific metabolic rate increases more slowly. However, counterintuitively, the measure of worker sensitivity to changes in temperature, as assessed by the instantaneous Q10 value (i.e., the rate of change across 10°C temperature intervals), shows that large ants are more sensitive than small ants to changes in temperature in terms of both water loss and metabolic rate. Such differential thermal sensitivity allows to make testable predictions on the temporal distribution of foraging activity among workers of different sizes in polymorphic ant species, as well as how these species may alter their colony demographics in response to rising temperatures.

## 1. INTRODUCTION

Body size is a factor of paramount importance at all levels of biological organization (Schmidt-Nielsen 1984). It can impact not only the morphology, anatomy, behavior and physiology of organisms, but also their evolution and ecology through its implication in trophic interactions (La Barbera 1989, Kalinkat et al. 2015). The link between body size and physiology is particularly strong in ectotherms because body size determines how quickly they gain or lose heat from external sources. Although their metabolic rate, and therefore their level of activity and their capacity to interact with the environment, depends on a variety of internal and external factors, it is essentially determined by the tight interaction between ambient temperature and body size (Glazier & Gjoni 2024).

Because of their small size insects are characterized by a low thermal inertia. Consequently, their body can warm up very quickly with rising temperatures, leading to greater energy expenditure and to potentially fatal risks of dehydration. The study of the interaction between body size, temperature and physiology in insects is best achieved in the adult stage of holometabolous insects. In fact, at this stage none of the anatomical and physiological changes associated with developmental processes in the pre-adult stages of heterometabolous insects can interfere with their thermal physiology. In this regard, ants make a very good model for the study of within-species interaction between body size and temperature because, in addition to morphological differences between individuals of the reproductive and non-reproductive caste, approximately 13% of ant species also exhibit a polymorphism of the worker caste, i.e. a variation in the size and/or shape of individuals (Hölldobler & Wilson 1990, Fjerdingstad & Crozier 2006). This polymorphism is generally closely associated with division of labor within colonies (Wills et al. 2018), one of the features that has largely contributed to the worldwide ecological success of social insects (Taborsky et al. 2025). In some ant species the worker caste can be further divided in several distinct physical subcastes and a subcaste of large workers, called soldiers, generally in charge of the defense of the colony, can be identified (Hölldobler & Wilson 1990).

The large variation in body sizes observed in the worker caste of polymorphic ant species is not only associated with differences in their morphology and behavior but also with differences in their internal anatomy, e.g. their brain (Kelber et al. 2010, Muratore et al. 2022) and muscle (Khalife et al. 2022) structure and organization, as well as their sensory physiology (vision: Narendra et al. 2011, Arganda et al. 2020, olfaction: Lopez-Riquelme et al. 2006, Kleineidam et al. 2007, Gordon et al. 2018, Ferguson et al. 2023, gustation: Josens et al. 2018). However, the main impact of body size in ant workers lies in their capacity to react to ambient temperature, which depends on their thermal sensitivity. In species of ants characterized by worker polymorphism, the general rule is that large workers are more resistant to heat than small workers (leaf-cutting ants: Ribeiro et al. 2012, Bouchebti et al. 2015, Baudier & O’Donnell 2019, Welch et al. 2020; seed harvesting ants: O’Donnell et al. 2020, Arnan et al. 2022; army ants: Baudier et al. 2015, Baudier & O’Donnell 2018 ; *Cataglyphis*: Cerda & Retana 1997, 2000, Clémencet et al. 2010, Wendt & Verble-Pearson, 2016). This resistance is usually measured by the critical thermal maximum (CTmax), i.e. the temperature at which knockdown or the onset of muscle spasms is observed. However, CTmax is a measure of mortality at extreme temperatures and is thus a rough measure of the thermal tolerance of individuals. Moreover, there are some methodological issues because the evaluation of CTmax may vary according to the procedure used, notably the duration of the experiment and the heating rate that is used to reach the critical temperature (Ribeiro et al. 2012). A much better approach is the measure of the thermal reaction norm of individual ants, i.e. the pattern of variation of their physiological response to increasing temperatures. This latter can be assessed by the measure of their metabolic rate (MR), which is directly linked to the amount of energy an organism uses to keep its body functioning. A general rule in insects (Lighton et al. 1987; Full & Tu 1991), as in other animals (Schmidt-Nielsen 1984), is that the mass-specific MR, i.e. the MR expressed per unit body mass, decreases with increasing body mass, meaning that small organisms consume more energy per unit body mass than large ones.

The objective of our study was to assess the effect of both temperature and body mass, taken as a proxy of body size, on general activity, water loss and MR in the Mediterranean polymorphic seed harvesting ant *Messor barbarus*. The colonies of this species can contain several tens of thousands of individuals (Cerdan 1989) and are characterized by a strong polymorphism of the worker caste, with body mass varying between 1.5 to 40.0mg (Bernadou et al., 2016). This polymorphism is characterized by a continuous monophasic allometry between head mass and body mass (Merienne et al. 2020), so that large workers have proportionally a bigger head compared to small workers. First, we hypothesize that, because small ants have a higher body surface area to body mass ratio, water loss for a given temperature should be higher in small ants than in large ants. In addition, since large workers have a more sclerotized head capsule than small ones, which could reduce cuticular water loss, we expected that water loss in small ants should increase more rapidly than in large ants. Second, we hypothesize that the negative relationship between mass-specific MR and body mass observed in individuals of different sizes belonging to different species should hold for intraspecific comparison. Therefore, we expected that, for a given temperature, large workers should consume less energy per unit body mass than small ones and thus that the regression curves of mass-specific MR against temperature should have a higher elevation for small than for large ants. Finally, since MR depends essentially on body size and temperature, we expected that the mass-specific MR should increase in the same way with temperature in ants of different sizes.

## 2. METHODS

### (a) Studied species and rearing conditions

We worked on three colonies of *Messor barbarus* collected in southern France (42.802, 2.976) on the French Mediterranean Coast. Each colony was housed in the laboratory in a plastic box which contained several test tubes wrapped with white paper and partially filled with water retained by a cotton plug in which the ants established their nest. The colonies were kept in a room at a temperature of 25°C with 50% relative humidity. They had access to water *ad libitum* and were fed daily with a mixture of seeds of various species.

### (b) Mass-specific Metabolic rate (msMR) measurements

The MR of individual workers was measured at five different temperatures (15, 20, 25, 30, and 35°C) with a high resolution respirometric system (Sable Systems Europe GmbH, Berlin, Germany, https://www.sablesys.com/) equipped with a nondispersive infrared CO_2_ analyzer (LI-COR 850, LI-COR, Lincoln, NB, USA, https://www.licor.com/, lower limit detection: 1.5ppm). The temperatures chosen lie in the range of soil surface temperatures for which a high level of foraging activity is observed in *M. barbarus* in its natural environment (Azcarate et al. 2007, Arnan et al. 2011). During the trials, ants were placed individually in seven cylindrical glass respirometric chambers (diameter: 2cm, length: 7cm, effective air volume: 14.14cm^3^) housed in an insulated box (PELT-DROP-IN) whose temperature was continuously recorded with a thermistor cable and regulated by a Peltier controller (PELT-5, control stability: 0.01-0.20°C). An eighth chamber remained empty and was used for baselining. The inside of the box was lighted by LEDs (Cineroid L10-BC).

We performed constant volume respirometry (Lighton & Halsey 2011). In each trial ants were kept in the airproof respirometric chambers for a total of three hours. At 30min intervals the air of each chamber was flushed for 200s and replaced by fresh air scrubbed of H_2_O and CO_2_ through the passage in a drierite/ascarite/drierite column. The flow rate was set at 50 ml/min by a subsampler pump (SS-4) that was regulated by an Alicat Scientific MC Series valve (MFCV-31) connected to a mass flow control unit (MFC-2). The air from the chambers first passed through a column of magnesium perchlorate (Cl_2_MgO_8_) to remove the water vapor due to the insects’ respiration and then was directed to the CO_2_ analyzer to measure the CO_2_ (expressed in μl CO_2_ hr^−1^) accumulated in the chambers. A multiplexer (RM-8) programmed by a software provided by Sable Systems (SW-EXPEDATA-P) operating through an interface unit (UI-3), allowed to flush the air sequentially from the eight respirometric chambers. The MR measurements were obtained automatically by the software through the integration of the area below the CO_2_ peaks of the excurrent air flow. The CO_2_ values measured from the empty chamber were subtracted from the values obtained from the chambers containing ants to correct for possible leaks. Since there was little variation in the measure of MR over the five 30-min sessions (Fig. S1), the MR values measured for each ant were averaged over the last five 30-min sessions of the trials. Indeed, since the air was not scrubbed of CO_2_ at the moment ants were installed in the chambers, the MR measurements of the first 30min session were excluded from the analysis. The CO_2_ analyzer was calibrated every three respirometric trials, zeroing it with N_2_ gas and then spanning it with a gas of known CO_2_ concentration (5,000 p.p.m. CO_2_ in N_2_ ± 1%).

Ants from three different colonies of *M. barbarus* were used in the trials. For each experiment seven ants chosen randomly were captured directly in the foraging box of the colonies. The size of the ants was assessed visually and categorized in three size classes as small, medium or large. Ants were then weighed a first time with an analytical balance (Metler Toledo MS105: accuracy: 0.1mg) to obtain their fresh mass. This fresh mass was used to calculate the mass-specific MR (msMR in μlCO_2_.h^−1^.mg^−1^), i.e. the value of MR obtained from the SW-EXPEDATA-P software divided by the fresh mass of the ants. The ants were then placed in the respirometric chambers. To avoid any perturbation due to pheromone deposits, the respirometric chambers were washed with a neutral detergent and dried with paper towel every time ants from a different colony were used. All ants were immediately weighed at the end of the experiment to assess the mass they lost by evaporation during the experiment. They were then placed in an oven at 60°C during 24h to measure their dry mass. In total, the msMR of 144 ants was measured for each of the five temperatures tested.

### (c) Percentage of water loss (PWL) measurements

The percentage of body water lost by the ants at the 5 temperatures tested during the 3 hours they spent in the respirometric chambers at 0%RH was determined gravimetrically. Following Arnan & Blühtgen (2015), we used the following equation to calculate percent water loss:

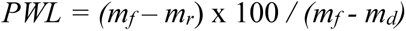

where *m_f_* is the fresh mass (mg) of the ants before being placed in the respirometric chambers, *m_r_* is their mass (mg) after the 3 hour-respirometer session and *m_d_* is their dry mass (mg).

### (d) Ant activity measurements

To measure ant activity during the respirometer trials, a webcam (Logitech Brio 4k ultra-HD), fixed on the inner side of the cover of the insulated box in which the respirometric chambers were placed, filmed the ants from above at a rate of 25fps. Ant activity in each respirometric chamber was then measured by an in-house software which calculates an activity score equal to the sum of the number of pixels that changes between two successive images for each 30min session of the respirometric trials. To correct for the different size of the ants, the activity score was divided by body mass raised to the power 2/3= 0.66 (Lighton & Feener 1989, Schilman et al. 2007) which gives a value proportional to body surface.

### (e) Data analysis

Water loss in insects occurs through both respiration and evaporation through the cuticle, with the latter representing more than 80% of water losses in many taxa (Chown 2002), including ants (Johnson et al. 2011). From a theoretical point of view, one should thus expect that the proportion of water lost by ants during the 3 hours they spent in the respirometric chambers should be proportional to their body surface area. Since this area increases as the square of body length whereas body mass increases as the cube, one should expect that the percentage of water loss would decrease allometrically with dry body mass with a 2/3 exponent, following the equation *PWL* ≈ *m_d_*^−0.66^ (Kühsel et al. 2017), which can be linearized as log(*PWL*) ≈ −0.66 log(*m_d_*). Consequently, the effect of body mass (mg) and temperature (°C) on *PWL* was tested with a GLM with dry body mass and *PWL* expressed on a log scale. In addition, to account for the fact that *PWL* may vary differently with body size with increasing temperatures, e.g. because of different thickness of the cuticle (Peeters et al., 2016), we also considered the interaction between body mass and temperature in the statistical model.

The influence of temperature and body mass on ants’ activity was tested with a GLM with a Gaussian error distribution. The interaction between body mass and temperature was included in the model because we hypothesized that activity may vary differently with temperature in ants of different sizes (Hurbert et al. 2008; Welch et al., 2020). To account for the fact that ant activity could reach a plateau at high temperatures we entered a quadratic term for temperature in the model.

The effect of body mass (mg) and temperature (°C) on the msMR (μlCO_2_.h^−1^.mg^−1^) in our experiment was tested with a GLM with both msMR and body mass expressed on a log scale. Since msMR may decrease non-linearly with increasing temperature (Shik et al., 2019), we entered a quadratic term for temperature in the model. Finally, since the effect of temperature on msMR may vary in ants of different sizes, we also considered the interaction between body mass and temperature in the statistical model.

According to the literature on animal physiology, the mass-specific Standard Metabolic Rate (msSMR), i.e. the mass-specific metabolic rate at rest also called basal metabolic rate, is higher in small animals than in big animals and scales allometrically with their body mass with a negative exponent varying between −0.25 and −0.33, corresponding to the surface area theory and resource transport theory, respectively (Glazier, 2018). To test whether this is also true with our ants, we assessed the msSMR of the ants by regressing the msMR corrected for activity on body mass and temperature with both msMR and body mass expressed on a log scale. For the same reason mentioned above, we entered a quadratic term for temperature in the model and considered the interaction between body mass and temperature.

Except for *PWL* the value of body mass used in the models was always the fresh mass of the ants. Moreover, temperature was considered as a continuous variable in all models. The value considered for each ant in the analyses of the msMR and the activity score was their average value over the last five 30-min sessions of each respirometric session. Note that, for technical reasons, the dry weight (and thus the *PWL*) could not be obtained for 10 ants (1.4% of total) and the activity score (and thus the msSMR) for 94 ants (13% of total).

To compare the thermal sensitivity of ants of different sizes as regards to the different variables measured (*PWL*, activity, msMR, msSMR) we calculated the Q_10_ value, which is commonly used in thermal physiology to study the reaction norm of animals across a temperature gradient (Shik et al., 2019). The Q_10_ corresponds to the rate of change in the variable measured across 10°C temperature intervals. To calculate the instantaneous Q_10_, we used the derivative of the equation of the statistical models that relate the value of the variables measured in each ant as a function of their body mass and temperature (Lighton & Bartholomew 1988, Vogt & Appel 1999; Lighton 2008; Shik et al. 2019, see SI S1).

For each variable, we started with the full model with interaction and then compared the nested models through AIC using the *MuMIn* R package (Barton 2025). When several models were equivalent we chose the simplest one. Model diagnostic was achieved with the function *check_model()* of the performance R package (Lüdecke et al. 2021). The impact of each independent variable in the model was assessed by examining the value of the model coefficients after their standardization using the *ggstats* R package (Larmarange 2025). All data analyses were run and figures generated with R4.5.0 run under RStudio2024.12.0.

## 3. RESULTS

The distribution of the fresh mass of the workers used in the experiment shows three size classes (Fig. S2): small or *minor* (≤ 5mg), *medium* (>5mg and ≤ 15mg), and large or *major* (>15mg) workers. The mass of the workers ranged from 0.70mg to 41.50mg; the median mass of each size class was 2.30, 9.40, 26.10mg for *minor*, *media* and *major* ants respectively.

### (a) Effect of temperature and body mass on water loss

There was a weak but significant interaction effect between temperature and body mass on the percentage of water loss (Table 1). The percentage of water loss increased slightly more rapidly with increasing temperature in small ants than in medium and large ants (Fig. 1a-b). Overall, the percentage of water loss increased non-linearly with increasing temperature and decreased allometrically with increasing body mass, with an exponent comprised in the interval [−0.870 −0.624] (Table 1), which thus includes the theoretical value of −0.66 corresponding to the ratio of body surface area to body mass. Body mass and temperature had an equivalent impact on the percentage of water loss (Fig. S3). The sensitivity of ants to a change in temperature in terms of water loss increased with increasing body mass (Fig. 1c). The instantaneous Q_10_ value of large ants was higher than that of medium ants which was itself higher than that of small ants (Fig. 1d). However, this sensitivity did not vary with temperature.

**Figure 1:**
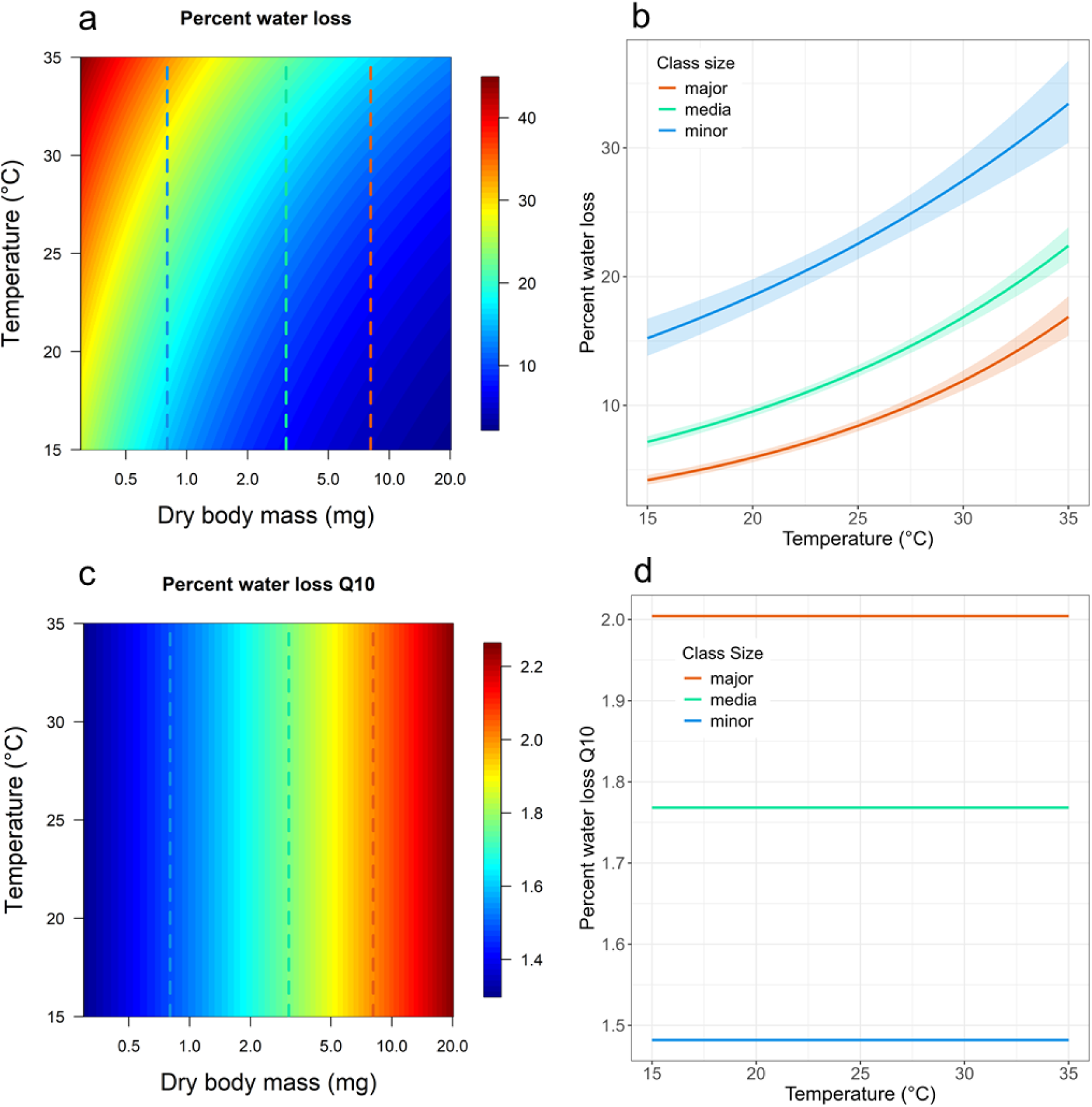
Percentage of water loss in *Messor barbarus* ants during the 3 hours spent in the respirometric chambers at 0%RH. (a) Percentage of Water Loss as a function of dry body mass and temperature. (b) Percentage of water loss as a function of temperature for the median dry mass of each worker size class. (c) Instantaneous Q_10_ values for Percentage of Water Loss as a function of dry body mass and temperature. (d) Instantaneous Q_10_ values for Percentage of Water Loss as a function of temperature for the median dry mass of each worker size class. The dashed vertical lines in (a) and (c) show the median values of the mass of the workers of each size class. *N*= 710 ants.

**Table 1.** Results of the statistical models used to investigate the effect of dry body mass (*m_d_*) and temperature on percentage of water lost (*PWL*) and of fresh mass (*m_f_*) and temperature on mass-specific MR (msMR) and mass-specific standard MR (msSMR) of *Messor barbarus* ants.

| Response variable | Factors | Coeff [CI<sub>95%</sub>] | SE | t | P | R<sup>2</sup> adj. |
| --- | --- | --- | --- | --- | --- | --- |
| log($PWL$) | log($m_d$) | -0.747 [-0.870 -0.624] | 0.062 | -11.921 | <0.001 | 0.62 |
|  | Temperature | 0.027 [ 0.016 0.039] | 0.005 | 4.800 | <0.001 |  |
| | log($m_d$) x Temperature | 0.013 [ 0.008 0.018] | 0.002 | 5.342 | <0.001 | |
| log(activity) | log($m_f$) | -0.650 [-0.674 -0.625] | 0.018 | -39.533 | <0.001 | 0.82 |
|  | Temperature | -0.084 [-0.114 -0.053] | 0.016 | -5.317 | <0.001 |  |
|  | Temperature<sup>2</sup> | 0.002 [0.001 0.003] | 0.000 | 6.024 | <0.001 |  |
| log(msMR) | log($m_f$) | -0.656 [-0.722 -0.590] | 0.034 | -19.490 | <0.001 | 0.82 |
|  | Temperature | 0.092 [0.069 0.116] | 0.012 | 7.736 | <0.001 |  |
|  | Temperature<sup>2</sup> | -0.001 [-0.002 -0.001] | 0.000 | -5.234 | <0.001 |  |
| | log($m_f$) x Temperature | 0.008 [0.006 0.011] | 0.001 | 6.256 | <0.001 | |
| log(msMR) | log(Activity) | 0.608 [0.560 0.655] | 0.024 | 25.07 | <0.001 | 0.51 |
| log(msSMR) | log($m_f$) | -0.268 [-0.355 -0.181] | 0.044 | -6.073 | <0.001 | 0.54 |
|  | Temperature | 0.136 [0.106 0.167] | 0.015 | 8.753 | <0.001 |  |
|  | Temperature<sup>2</sup> | -0.002 [-0.003 -0.002] | -0.000 | -7.292 | <0.001 |  |
| | log($m_f$) x Temperature | 0.008 [0.005 0.012] | 0.001 | 4.903 | <0.001 | |

### (b) Effect of temperature and body mass on activity

Ant activity increased with decreasing body mass (Table 1, Fig. 2) and increased non-linearly with increasing temperatures, in the same way for ants of different sizes. Temperature had a higher impact on ant activity than body mass (Fig. S4). The sensitivity of ants to a change in temperature regarding activity increased in the same way with increasing temperatures for ants of different sizes (Table 1, Fig. 2c-d): the higher the temperature, the higher the effect of a temperature increase on ant activity.

**Figure 2:**
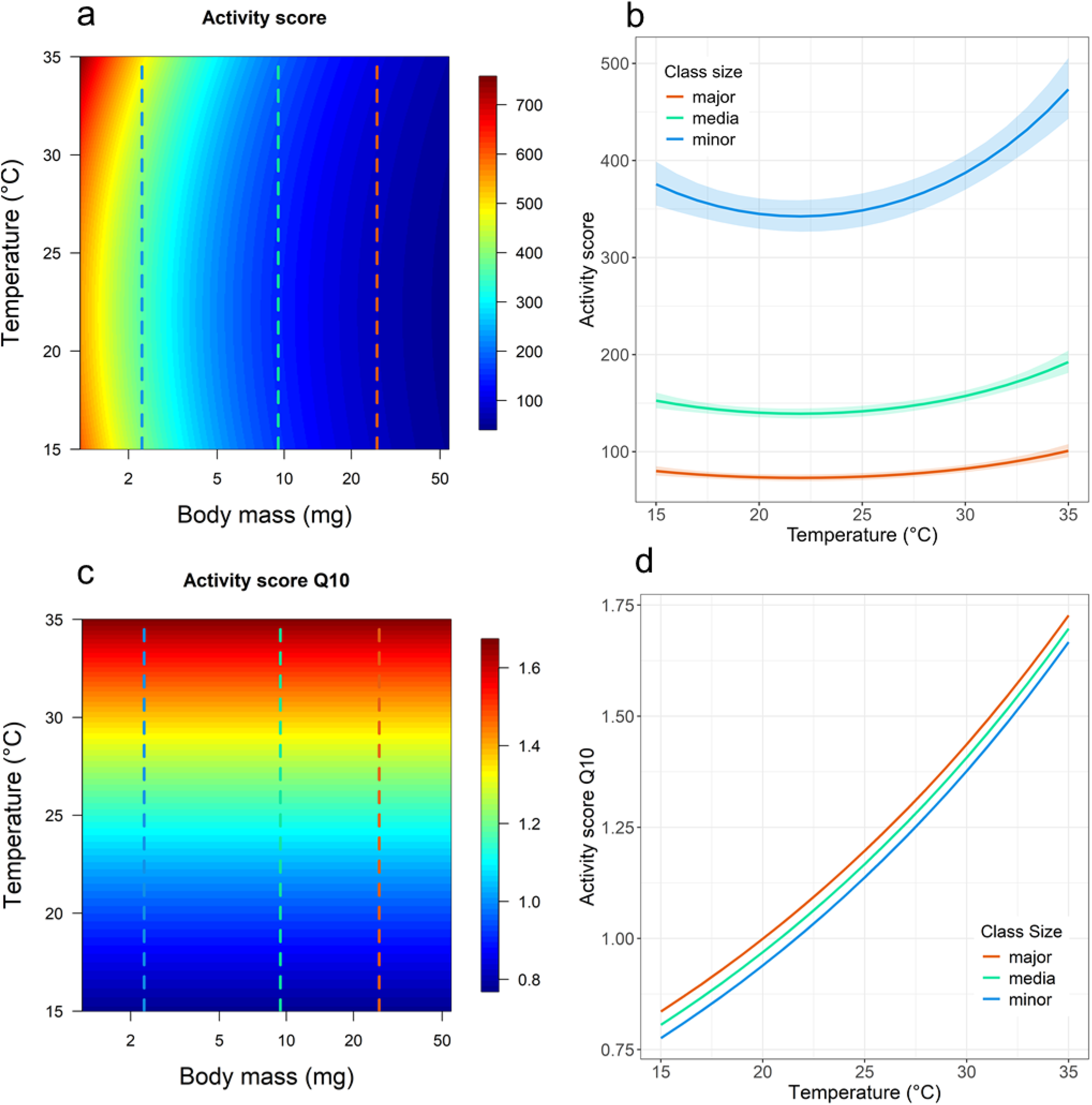
Activity score of *Messor barbarus* ants during the 3 hours spent in the respirometric chambers at 0%RH. (a) Activity score as a function of fresh body mass and temperature. (b) Activity score as a function of temperature for the median mass of each worker size class. (c) Instantaneous Q_10_ values for activity score as a function of body mass and temperature. (d) Instantaneous Q_10_ values for activity score as a function of temperature for the median mass of each worker size class. The dashed vertical lines in (a) and (c) show the median values of the mass of the workers of each size class. *N*= 626 ants

### (c) Effect of temperature and body mass on mass-specific metabolic rate

The msMR of ants varied slightly differently with temperature for workers of different sizes (Table 1). The msMR of small ants increased more rapidly with increasing temperature than that of medium or large ants (Fig. 3a-b) and was much more impacted by temperature than by body size (Fig. S5). The msMR instantaneous Q_10_ decreased with increasing temperatures and, whatever the temperature, was higher for large ants than for medium or small ants (Fig. 3c-d), showing that, in terms of the energy required to maintain their body functioning, large ants were more sensitive to a change in temperature than small ants.

**Figure 3.**
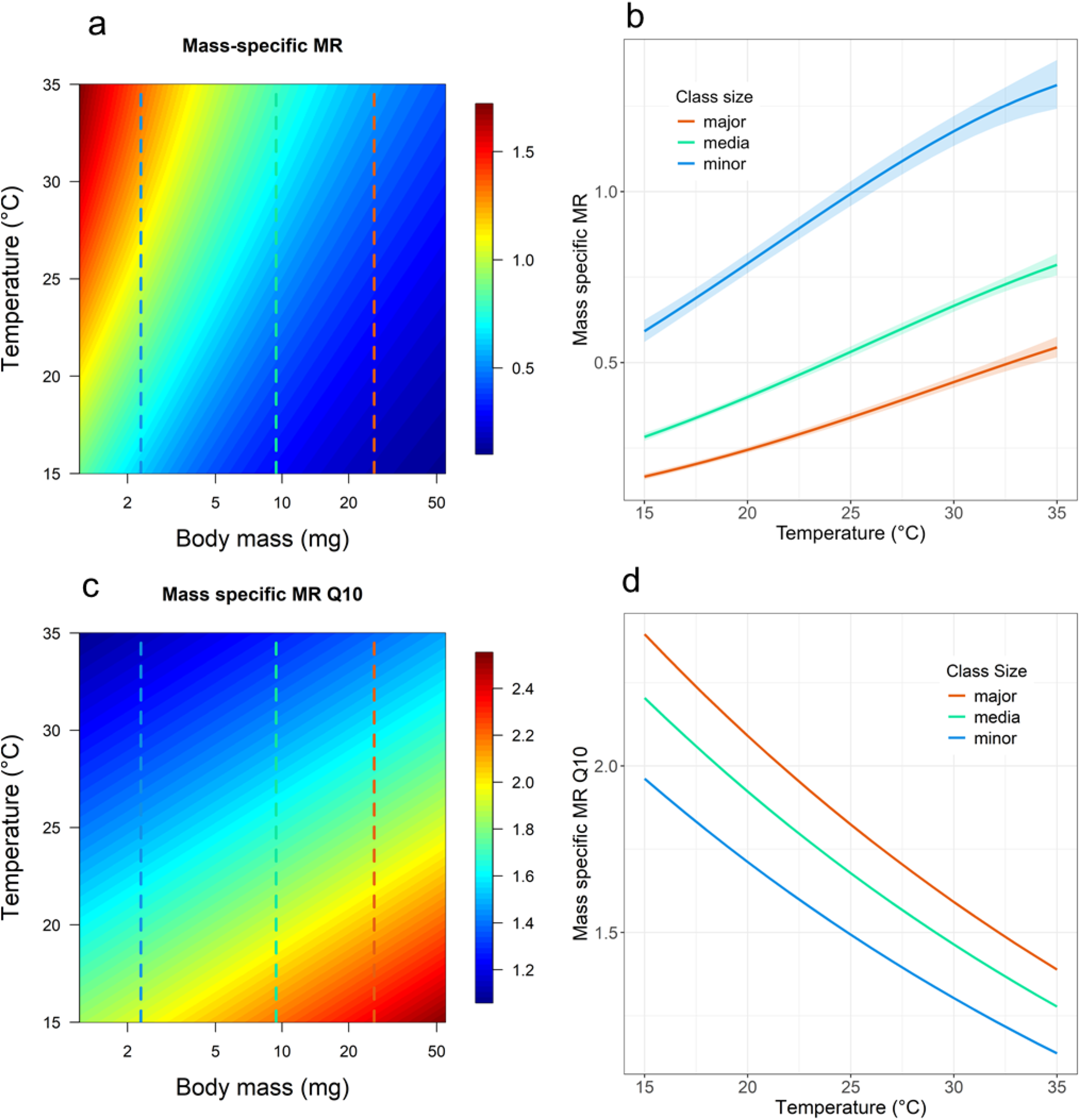
Mass-specific metabolic rate (in µlCO2.mg^−1^.h^−1^) of *Messor barbarus* ants. (a) Mass specifc MR as a function of fresh body mass and temperature. (b) Mass-specific MR as a function of temperature for the median mass of each worker size class (c) Instantaneous Q_10_ values of mass specific MR as a function of temperature and body mass. (d) Instantaneous Q_10_ values of mass specific MR as a function of temperature for the median mass of each worker size class. The dashed vertical lines in (a) and (c) show the median values of the mass of the workers of each size class. *N*= 720 ants.

### (d) Effect of temperature and body mass on mass-specific standard metabolic rate

As expected, the msMR increased significantly with increasing activity (Table 1). The msSMR, i.e. the msMR corrected for activity, varied in the same way as the msMR (Table1, Fig. S6) and, in the same manner as for msMR, temperature had the highest impact on msSMR (Fig. S7). The msSMR decreased allometrically with increasing body mass, with an exponent comprised in the interval [−0.355 −0.181] (Table 1), which thus includes both the theoretical values of −0.25 and −0.33 predicted by the surface area theory and resource transport theory, respectively.

## 4. DISCUSSION

Caste polymorphism is one of the main characteristics of social insects and is in part responsible for their ecological success (Taborsky et al. 2025). In ants for example, size variation within the worker caste has been shown to be important to keep a high level of foraging efficiency (Wilson 1983, Waser 1998, Reyes-Lopez & Fernandez-Hager 2001, Arnan et al. 2011, Constantino et al. 2021, Ramirez-Olier et al. 2022). In this study we found that large ants have a lower rate of water loss than small ants and, consequently, that they are more tolerant to desiccation, one of the most stressful factors in insects. In addition, we found that, depending on ant size, mass-specific MR is differently influenced by temperature: mass-specific MR increases slightly more rapidly with increasing temperatures in small ants than in large ones. However, calculation of the Q_10_ value, a measure of the sensitivity of ants to changes in temperature, shows that large ants are more sensitive than small ones regarding temperature variation: whatever the initial temperature, large ants show a stronger response than small ants to a change in temperature in terms of both water loss and metabolism. These results show therefore that ants of different sizes are characterized by different physiological strategies to cope with an increase in ambient temperature.

Our expectation was that large ants would lose proportionally less water with increasing temperatures than small ones because of their smaller body surface area to mass ratio (Lighton et al. 1994, Chown et al. 2011, Kühsel et al. 2017). Our results show that this was indeed the case: whatever the temperature, small ants had a higher percentage of water loss than large ants. Additionally, we found that small ants lost water more rapidly than large ants at increasing temperatures. Therefore, the percentage of water loss follows the same trend as the mass-specific MR (see below), which was predictable since respiratory water loss, linked to the exchanges of gas through the spiracles, is one of the two components, along with cuticular water loss, responsible for desiccation in insects (Chown 2002). In fact, as expected by other studies (Lighton & Bartholomew 1988, Shilman et al. 2005, Lighton & Turner 2008, Perl & Niven 2018), we found a positive relationship between the percentage of water loss and the msMR (Fig. S8).

In addition to body mass, differences in the physiology and/or morphology can also sometimes explain differences in the rate of desiccation of different sized workers. For example, differences in ventilation patterns (Lighton 1990, 1992, Perl & Niven 2018, Oladipupo et al. 2022), amounts of epicuticular lipids (Hood & Tschinkel 1990), cuticular hydrocarbon profiles (Gibbs 2002, Johnson & Gibbs 2004, Ajayi et al. 2020), and the thickness, pilosity, sculpturing (Buxton et al. 2021) or abrasion level (Johnson et al. 2011) of the cuticle are known to affect the rate at which ants lose water. In *M. barbarus,* large workers are not an enlarged copy of small ones because some of their body parts, their head in particular, increase allometrically with their body size (Bernadou et al. 2016). Therefore, large workers of *M. barbarus* may be more tolerant to high temperatures and more prepared to cope with heat than small ants, not only because of their size but also because of their larger head with its thicker cuticle.

In ants, as in all insects, locomotory speed increases with temperature (Hurlbert et al. 2008). Therefore, as could be expected, we found that ant activity inside the respirometric chambers increased with increasing temperature. What we did not expect however was that, for the same temperature, the relative activity of small ants would be higher than that of medium or large ants. Most field studies in ants show indeed that foraging activity decreases when hydric stress increases (Friedman et al. 2019). The higher activity observed in small ants in our experiment is likely to be related to the stress induced by the water loss they experienced during the respirometric trials. A positive relationship was indeed observed between activity and proportion of water loss (Fig. S9). Small ants may just increase their activity in an attempt to escape from a stressful situation.

As shown by other studies in ants (Traniello et al. 1985, Lighton et al. 1987, Bartholomew et al. 1988, Lighton & Bartholomew 1988, Lighton, 1989, Calabi & Porter 1989, Nilsen & Baroni-Urabni 1990, Weier et al. 1995, Lighton & Turner 2004, Chown et al. 2007, Clusella-Trullas et al 2010, Colinet et al. 2015, Kadochava et al. 2017, Perl & Niven 2018, Shik et al. 2019, Packard 2020), we found that, whatever the size of the workers, mass-specific MR was positively correlated with temperature. Mass-specific MR actually increased in a decelerating manner with increasing temperature. Although respiratory water loss is only a small portion of the overall water budget in ants (Lighton 1992), a reduction of metabolic activity could nonetheless be considered as a strategy to cope with high temperatures since it contributes to reducing water loss through respiration (Perl & Niven 2018). Moreover, as expected by the data from the literature, mass-specific MR was negatively correlated with body size at all temperatures. This may explain why in ant species showing a high size variation of the worker caste, such as leaf-cutting ants or seed-harvesting ants, medium and large ants are overrepresented on foraging trails compared to small ants (Wilson 1980, Heredia & Detrain 2005, Ramirez-Olier et al. 2022), insofar as they need less energy per unit body mass to keep their body functioning. At the same time, this allows to increase foraging efficiency since larger ants can carry bigger and heavier loads back to the nest. Since mass specific MR increases more rapidly with temperature in small ants than in big ants, one could also predict that the proportion of large ants on foraging trails should increase when ambient temperature increases. In a study of the foraging pattern of three sympatric species of *Messor,* Arnan et al. (2022) found indeed a slight, albeit non-significant, change in the distribution of forager sizes in *M. barbarus*: large workers were slightly more present on artificial baits shortly before foraging ended, when temperature was high, than at the start of the foraging period, when temperature was at least 10°C lower.

In the same way as the msMR, the msSMR, i.e. the metabolic rate corrected for activity, increased with increasing temperature and with decreasing body mass. Such relationships have been reported in a lot of animals, including ants (Riveros & Enquist 2011, Waters 2014, Perl & Niven 2018, Packard 2020). The allometric exponent of the mass-specific SMR against body mass, calculated across a variety of animal ranging in size between insects to elephants, has been found to be around −0.25 (Schmidt-Nielsen, 1984), which is the value expected under the nutrient supply network model of the metabolic theory of ecology (Chown et al. 2007). The alternative theory, that of the cell size model which states that the metabolic rate is a by-product of the way in which body size changes, predicts values for the allometric exponent around −0.33 (Chown et al. 2007). The value we found lies in between −0.355 and −0.181 and it is thus compatible with both theories and with all exponent values found in other species of ants so far (Fig. S10).

In terms of body mass, the sensitivity of ants to a change of temperature, calculated as an instantaneous Q_10_ value, followed the same trend for the percentage of water loss and the mass specific MR: although the percentage of water loss and mass-specific MR were lower for large ants than for small ones at all tested temperatures, large ants were more sensitive to changes in temperature. One might think that this higher sensitivity could be compensated for by large foraging ants walking faster to avoid exposure to rapid temperature changes but speed is not related to body size in *M. barbarus*. (Merienne et al. 2021). However, this may be offset by the fact that large ants have higher thermal inertia than small ants, meaning their internal temperature changes more slowly. (Kaspari et al. 2015). Whereas the sensitivity of ants in terms of water loss did not change with temperature, their sensitivity in terms of mass specific MR decreased with increasing temperatures, whatever their size. This result is concordant with that found in *Solenopsis invicta* (Vogt & Appel 1999) and in *Aphaenogaster senilis* collected above 1000m elevation in southern Spain (Shik et al. 2019) but at contrast with that found in the African ant *Camponotus fulvopilosus* for which the Q_10_ increases for temperatures above 25°C (Lighton 1989). Since sudden temperature increases from low temperatures are associated with higher energy expenditure in *M. barbarus*, one could predict that foraging workers should avoid environments where they are repeatedly exposed to large variations in ground surface temperature, e.g. areas with a patchwork of hot and cool spots linked to the absence or presence of vegetation coverage, respectively. This is actually the case as *M. barbarus* nests in grassland areas with low vegetation and avoids foraging in shaded areas (Azcarate & Peco 2003). Therefore, foraging workers probably experience small differences in soil temperature when foraging for seeds outside their nest. The critical event for them is thus the moment at which they leave their underground nest to emerge in the outdoor environment. However, in *M. barbarus,* the seeds are stored in shallow chambers, near the soil surface, in order to maintain a low moisture level and prevent germination (Cerdan 1989). Ants foraging for seeds and commuting between these chambers and seed patches should thus experience only a small difference in ambient temperature when they leave their nest. Our result on the mass-specific MR Q_10_ also shows that, whatever the initial temperature, large ants are more sensitive to a change in temperature than small and medium ants. Therefore, one could predict that large ants compared to small ants should decrease the frequency at which they commute between their nest and the outdoor environment. In fact, this is precisely what occurs in *M. barbarus* in which most of the seed transport is achieved by medium workers while large foraging workers are more specialized in cutting spikelets and therefore spend more time on seed patches.

In conclusion, measuring the thermal sensitivity of different-sized workers of *M. barbarus* in terms of water loss and metabolic rate allowed us to make testable hypotheses on the allocation of foraging tasks between different-sized workers and on the temporal distribution of their foraging activity. This approach could be used to better understand differences in worker size distribution, within as well as between species, in different thermal environments in ant species characterized by a polymorphism of the worker caste (Kaspari et al. 2015). It could also help predict how ant species might alter their colony demographics to adapt to the differential thermal sensitivity of their workers (Baudier & O’Donnell 2017) in response to rising temperatures in the context of global climate change (Arnan & Blüthgen 2015, Chick et al. 2017).

## Data availability

Dataset and R script to analyze the data and generate the figures of this paper are available on Zenodo (https://doi.org/10.5281/zenodo.17803760).

## SUPPLEMENTARY MATERIAL

### S1. Calculation of Q_10_ for water loss and mass-specific MR

#### S1.1. Q_10_ for Percentage of Water Loss

Equation of the model:

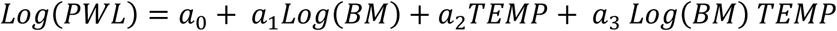

Derivative:

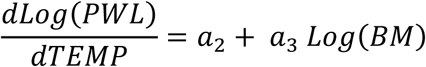

Q_10_:

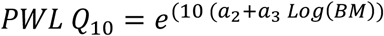

#### S1.2. Q_10_ for mass-specific MR

Equation of the model:

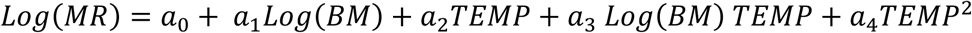

Derivative:

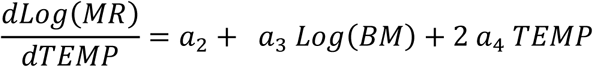

Q_10_:

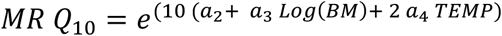

*MR*: Metabolic rate
*BM*: fresh body mass of the ants before being placed in the respirometric chambers
*TEMP*: temperature inside the respirometric chambers
*a_0_*: intercept of the model
*a_1_*: coefficient for fresh body mass
*a_2_*: coefficient for the temperature
*a_3_*: coefficient for the interaction between fresh body mass and temperature
*a_4_*: coefficient for the quadratic term of temperature

**Figure S1:**
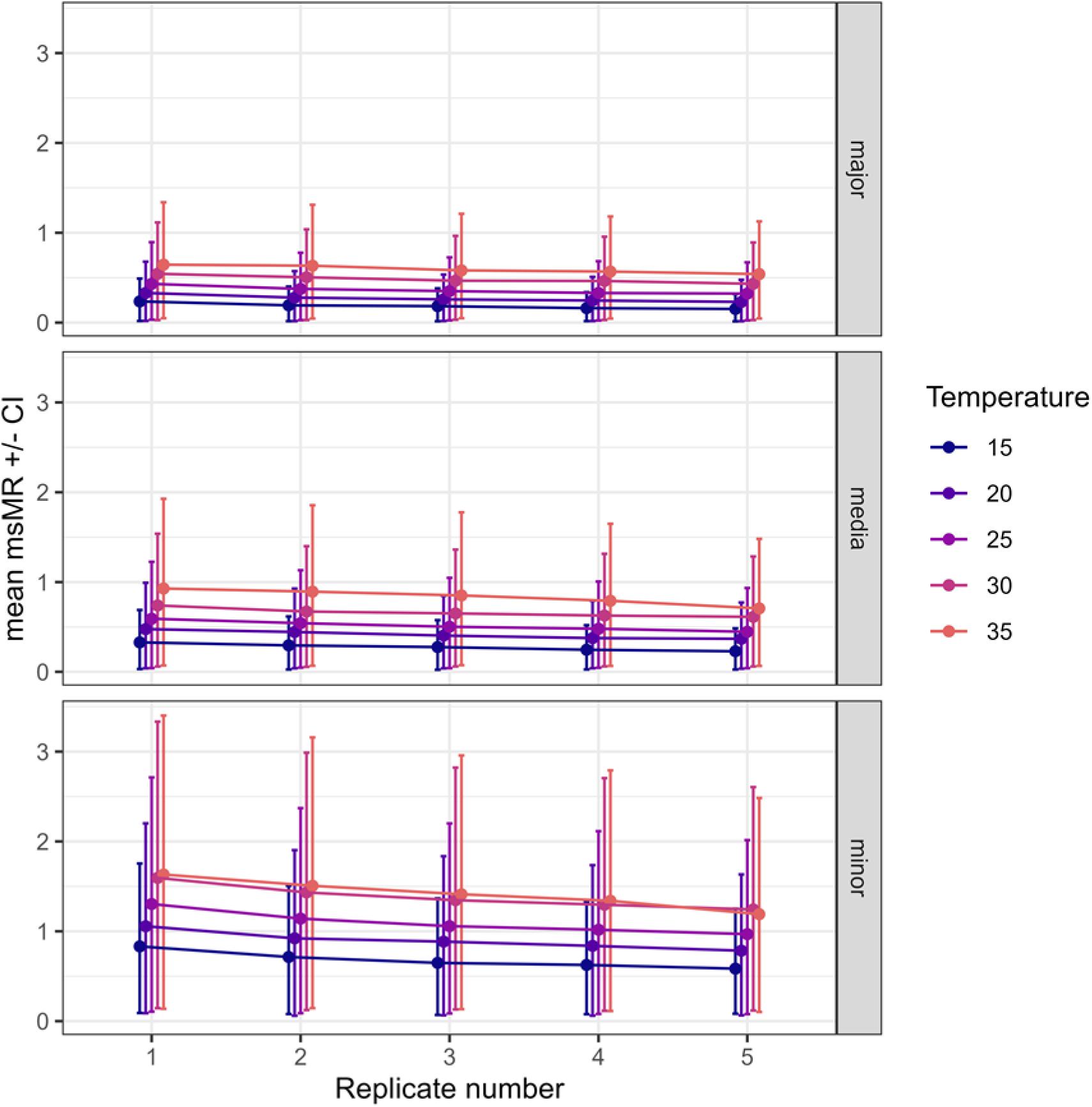
Change in mass-specific MR over the five successive replicates of the respirometric trials for each temperature and worker size class of *Messor barbarus* ants.

**Figure S2.**
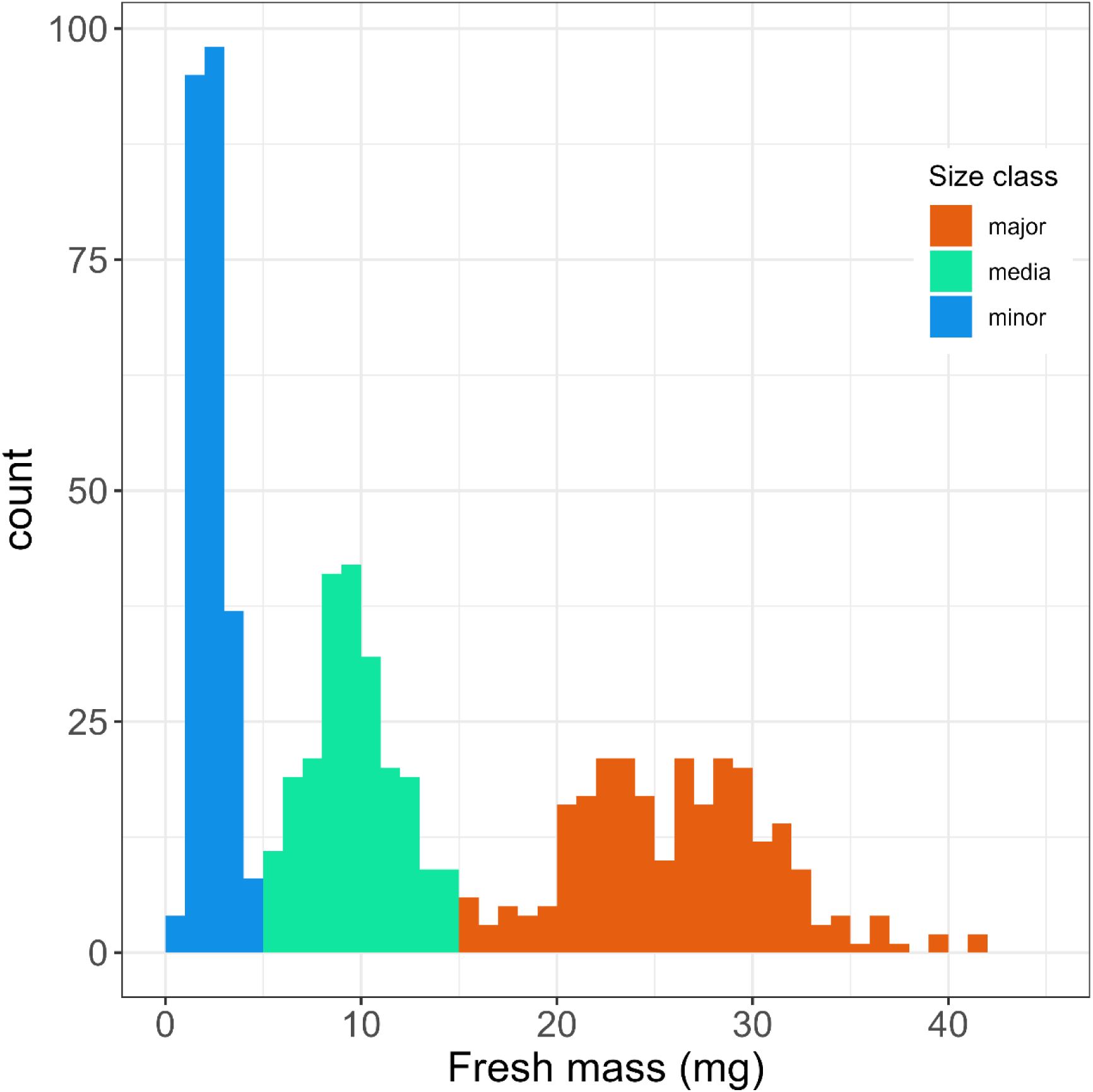
Frequency distribution of the fresh mass of *Messor barbarus* ants used in respirometric trials. N= 720 ants.

**Figure S3:**
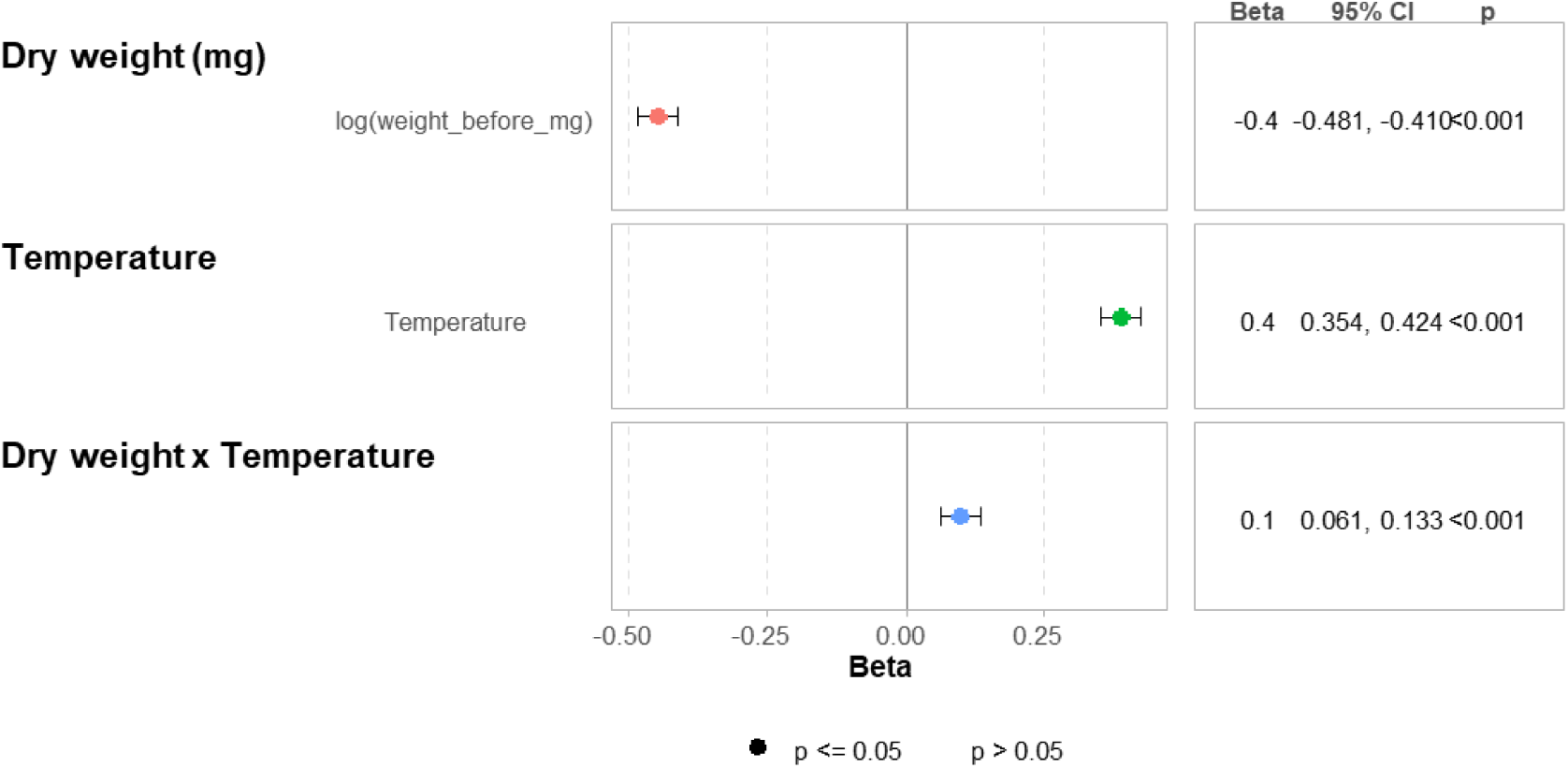
Standardized estimated coefficients with 95% confidence intervals of the statistical model of the Percentage of Water Loss as a function of dry body mass and temperature in *Messor barbarus* ants.

**Figure S4:**
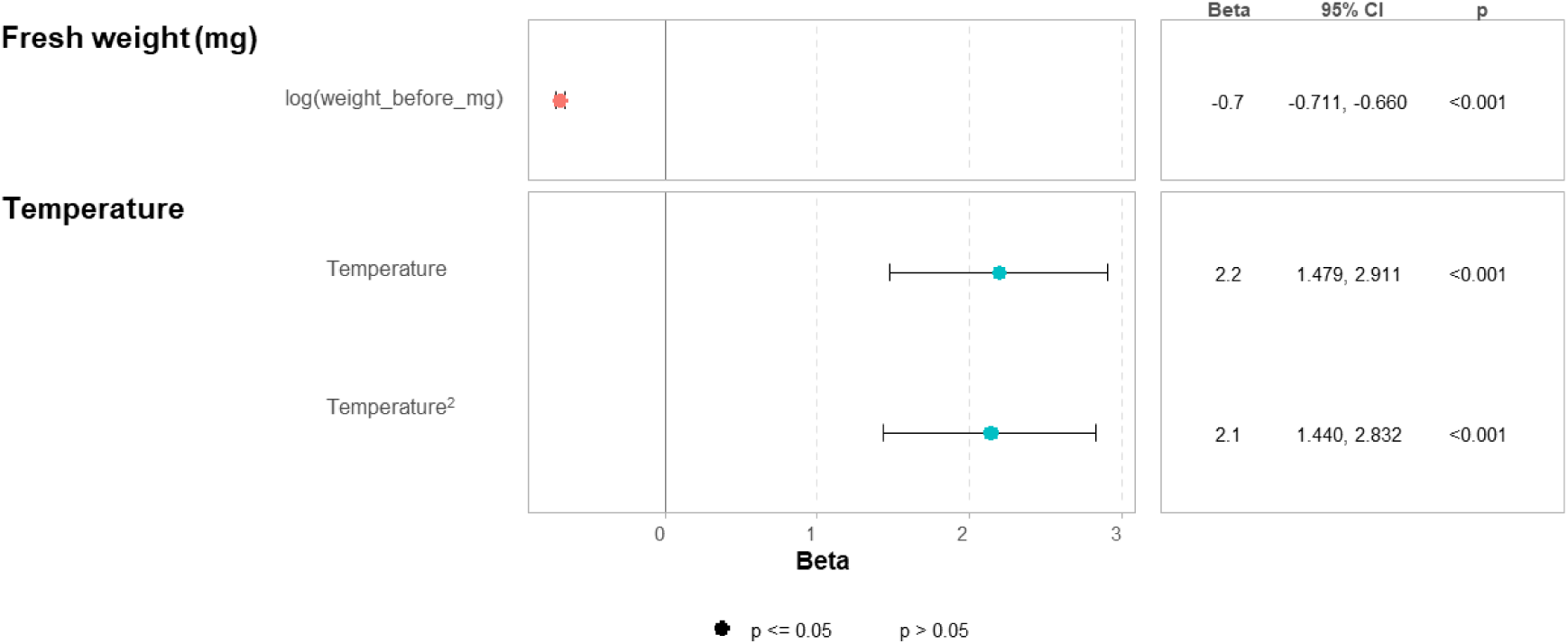
Standardized estimated coefficients with 95% confidence intervals of the statistical model of relative activity as a function of fresh body mass and temperature in *Messor barbarus* ants.

**Figure S5:**
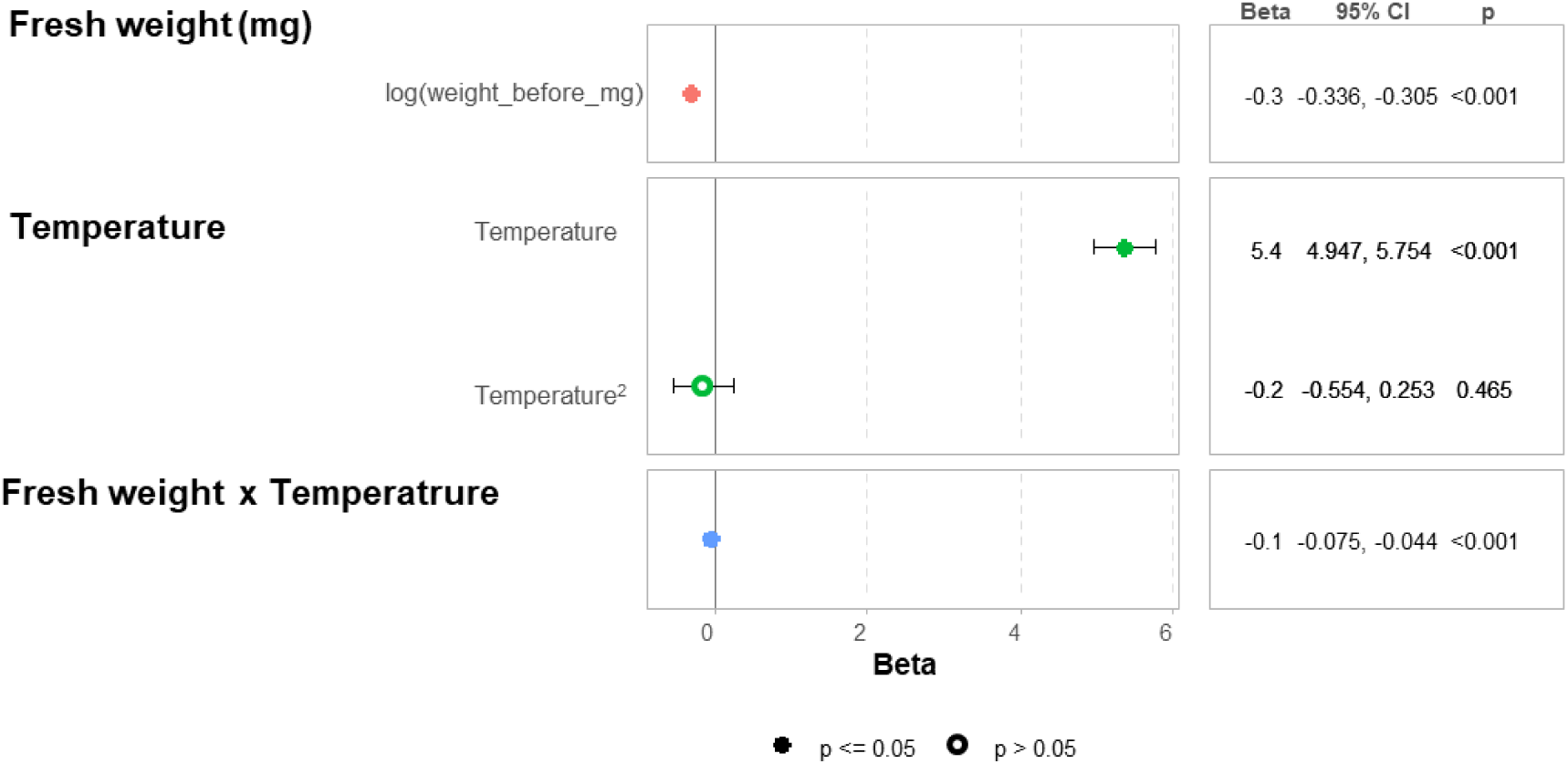
Standardized estimated coefficients with confidence intervals of the statistical model of mass-specific MR as a function of fresh body mass and temperature in *Messor barbarus* ants.

**Figure S6.**
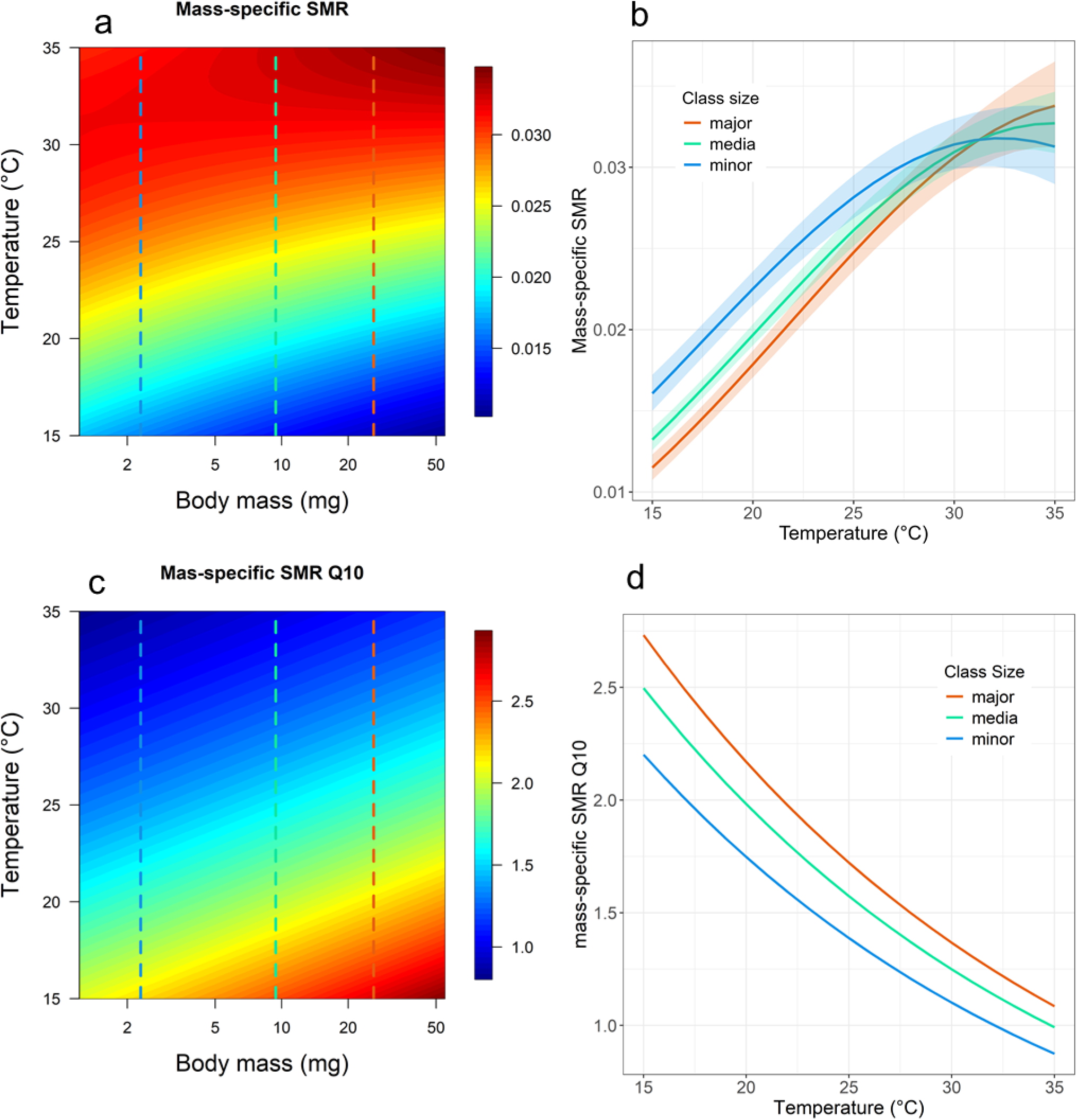
Mass-specific Standard Metabolic Rate (in µlCO2.mg-1.h^−1^) of *<u>Messor</u> <u>barbarus</u>* ants. (a) Mass specifc SMR as a function of fresh body mass and temperature. (b) Mass-specific SMR as a function of temperature for the median mass of each worker size class (c) Instantaneous Q10 values of mass specific SMR as a function of temperature and body mass. (d) Instantaneous Q10 values of mass specific SMR as a function of temperature for the median mass of each worker size class. The dashed vertical lines in (a) and (c) show the median values of the mass of the workers of each size class. N= 626 ants.

**Figure S7:**
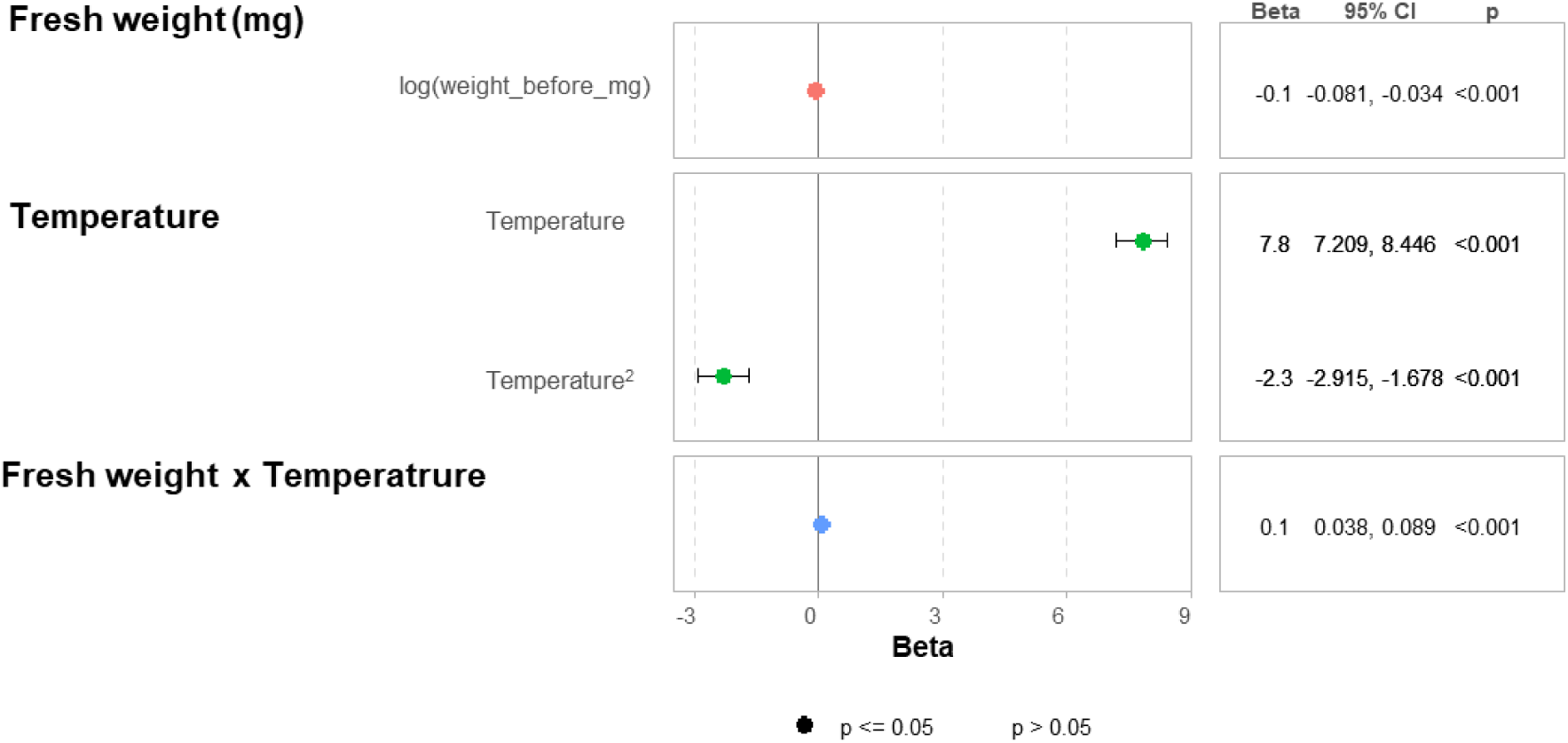
Standardized estimated coefficients with confidence intervals of the statistical model of mass-specific SMR as a function of fresh body mass and temperature in *Messor barbarus* ants.

**Figure S8:**
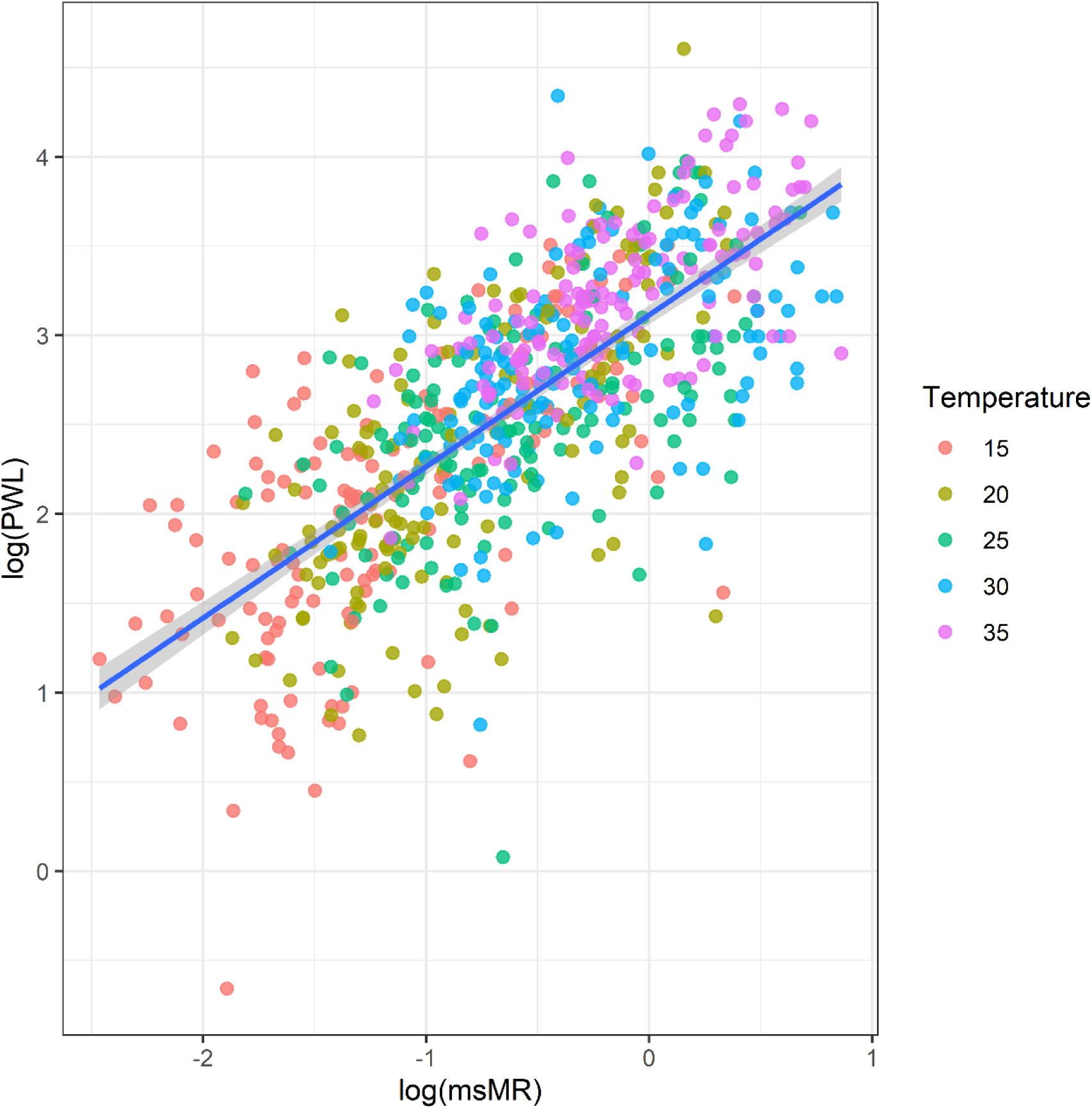
Percentage of Water Loss as a function of mass-specific MR in *Messor barbarus* ants. The equation of the regression line is: *log(PWL)= 3.115 + 0.849 log(mass-specific MR)*. R^2^= 0.521. The shaded area shows the confidence interval of the slope of the regression line.

**Figure S9:**
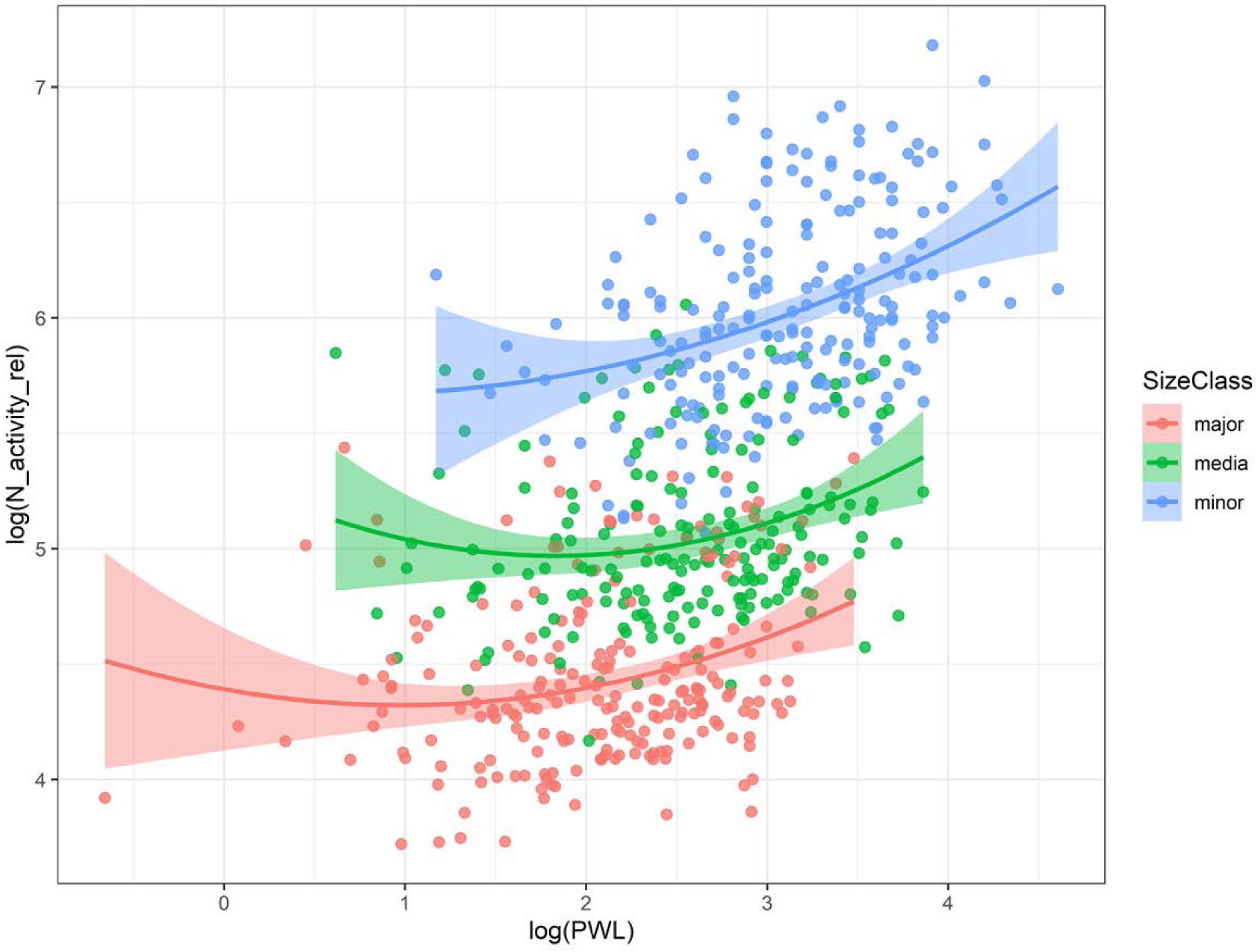
Ant activity score as a function of Percentage of Water loss in Messor barbarus ants. The equation of the regression lines are: *log(Activity)*= *4.45 – 0.16 log(PWL)* + *0.07 log(PWL)^2^, log(Activity)*= *5.00 – 0.16 log(PWL)* + *0.07 log(PWL)^2^, log(Activity)*= *5.84 – 0.16 log(PWL)* + *0.07 log(PWL)^2^*, for major, media and minor workers respectively. R^2^= 0.76.

**Figure S10:**
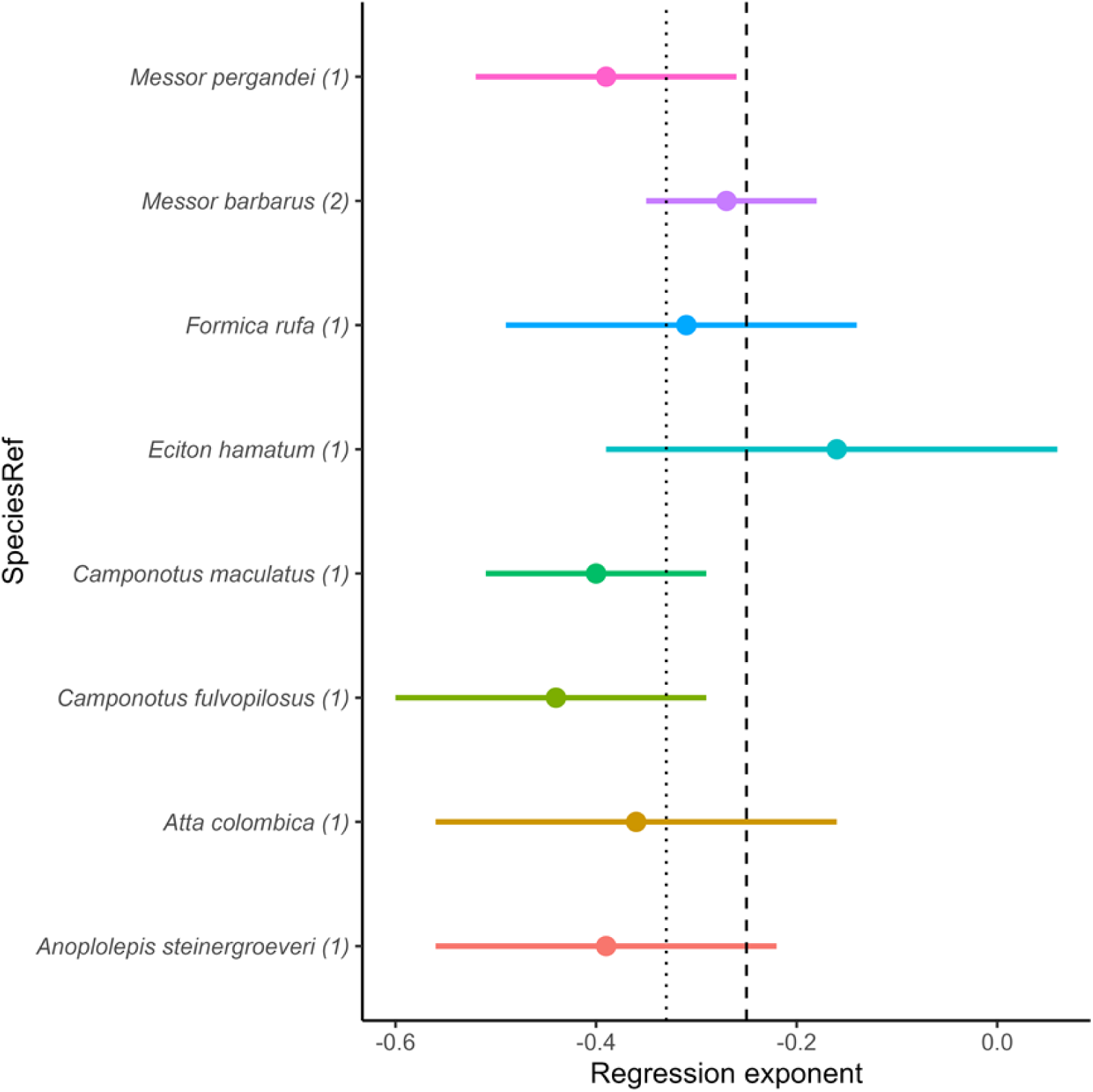
Values of regression exponent of mass-specific Standard Metabolic Rate with 95% confidence interval as a function of body mass for different species of ants. The dashed vertical line highlights the value −0.25 and the dotted line the value −0.33 expected under the surface area theory and resource transport theory, respectively. (1): Chown et al. 2007, (2): this study.

